# Neutrophils-Nitroblue Tetrazolium staining: A potential novel marker of women infertilely?

**DOI:** 10.64898/2026.08.02.742324

**Authors:** Maryouma M. Aghil, Abdulhakim S. Elnfati, Bashir A. Lwaleed

## Abstract

**Rationale:** Oxidative stress, resulting from an imbalance between the production of reactive oxygen species (ROS) and antioxidant defence mechanisms, disrupts cellular redox homeostasis and contributes to damage of biomolecules, including nucleic acids, proteins and lipids. Such disturbances can impair intracellular signalling pathways and have been implicated in the pathophysiology of female reproductive disorders. Nitroblue Tetrazolium (NBT) is a distinctive dye that assesses cellular redox activity in neutrophils through the detection of superoxide anion (O_2_^−^). However, the relationship between neutrophil-derived oxidative stress and women infertility remains to be fully elucidated. Aim of the study: We aimed to investigate the role of neutrophil oxidative stress in women infertility using a novel Nitroblue Tetrazolium (NBT) method developed in our laboratory.

**Materials and methods:** Blood and serum specimens were obtained from a total of 100 Libyan women, comprising healthy fertile women (n = 21; controls) and infertile women (n = 79). Superoxide anion (O_2_^−^) generation in neutrophils was assessed using the novel NBT method, while malondialdehyde (MDA), a marker of lipid peroxidation, was determined using the thiobarbituric acid reactive substances (TBARS) assay.

**Results:** A significant increase in both NBT-reactivity levels and the percentage of NBT-positive neutrophils was observed in infertile women compared with healthy fertile controls (P < 0.0001). In addition, MDA levels were significantly higher in infertile women than in the control group, indicating enhanced lipid peroxidation. MDA levels were positively correlated with NBT-reactivity levels (r = 0.410, P = 0.0001) and the percentage of NBT-positive neutrophils (r = 0.21, P = 0.047). A significant positive correlation was also observed between NBT-reactivity levels and the percentage of NBT-positive neutrophils (r = 0.510, P = 0.0001), demonstrating a close association between neutrophil oxidative activity and lipid peroxidation in women infertility.

**Conclusion:** The present study demonstrates the utility of a novel NBT method for detecting reactive oxygen species (ROS), a marker of oxidative stress, in neutrophils from both blood and serum specimens. The findings demonstrate increased neutrophil-derived oxidative stress in infertile women and suggest that this method may have potential as a diagnostic tool for the assessment of women infertility.

**Impact:** This study provides evidence that neutrophil-derived oxidative stress is significantly increased in women infertility and demonstrates the application of a novel Nitroblue Tetrazolium (NBT) assay for assessing oxidative stress in both blood and serum specimens. The significant associations between NBT-reactivity, NBT-positive neutrophils and malondialdehyde levels provide further insight into the contribution of oxidative stress to female reproductive pathophysiology. These findings support the potential utility of the novel NBT method as a simple and reliable diagnostic approach for evaluating oxidative stress in women infertility and provide a foundation for future studies investigating oxidative stress biomarkers in reproductive medicine.

## INTRODUCTION

Multiple enzymatic cellular mechanisms occur in the cytosol, mitochondria, endoplasmic reticulum, peroxisomes, and plasma membrane can produce reactive oxygen species (ROS). These molecules are short-lived, unstable, and highly reactive (Darbandi *et al*., 2018; Reczek & Chandel, 2015; Snezhkina *et al*., 2019). Mainly, the important sources of ROS in Graffian follicles are neutrophils, macrophages and granulosa cells through activation of steroidogenic cytochrome P450 enzyme, nicotinamide adenine dinucleotide phosphate (NADPH) oxidases (NOXs), xanthine oxidase, monoamine oxidases (Agarwal, *et al*., 2012; Lu *et al*., 2018; Snezhkina *et al*., 2019), and cyclooxygenase (Gupta *et al*., 2009; Pizzino *et al*., 2017).

During cellular mechanisms, oxygen (O_2_) is converted to superoxide anion (.O_2_^-^), which turns to hydrogen peroxide (H_2_O_2_) by superoxide dismutase (SOD) (Ahmad *et al*., 2017). However, when superoxide and hydrogen peroxide are not transformed, they act together to produce toxic hydroxyl free radical (. OH) (Darbandi *et al*., 2018; Snezhkina *et al*., 2019).

Furthermore, the production of free radicals can be enhanced exogenously by psychological stress, obesity, infection, and endocrine disrupting chemicals (EDCs) (Agarwal *et al*., 2012; Al-Gubory, 2014; Darbandi *et al*., 2018). The EDCs are ubiquitous chemicals and found in different products, most commonly in plastic bottles, toys, cosmetics, detergents, metal food cans and pesticides (Huo *et al*., 2015; Yang *et al*., 2015). The mechanism of EDCs is either by blocking or mimicking endocrine action, particularly, they can interfere with steroidogenesis, receptor binding and hormonal action (Rattan *et al*., 2017). The exogenous sources of ROS modulate the pathophysiology of the endocrine system by inducing oxidative stress and triggering redox-sensitive pathways, subsequently leading to inflammation and cell death. This may have irreversible effect on hypothalamic-pituitary-ovarian axis (Mahoney & Padmanabhan, 2010; Matuszczak *et al*., 2019).

The novelty of this work lies in using the NBT method to determine the presence of intracellular superoxide anion in neutrophils and serum of infertile women. The NBT is a pale yellow, water-soluble stain before being reduced by superoxide, which acts as electron acceptor. The principle of this method depends on the incubation of the sample with NBT in a particular condition to allow taking up NBT into endosomes (Volk and Moreland, 2014), where it interacts with superoxide anion causing NBT alteration and tetrazoinyl radical generation, tetrazole ring destruction and dismutation to form purple-blue insoluble crystals (Tvrda, 2019). Hence, tetrazolium salts are used widely to detect superoxide in neutrophils (Gosálvez *et al*., 2017). We hypothesized that NBT test could be a valuable marker of oxidative stress. Therefore, the aim of this study was to assess the positive response to NBT in the neutrophils obtained from blood and measuring O_2_^−^ in serum in fertile and infertile women.

## MATERIALS AND METHODS

### Study Population

The study was conducted on women recruited from Tripoli Center for Infertility Treatment, Tripoli, Libya. It consists of 100 women with an average reproductive age of (33.02 ± 5.44) years. Out of these, 79 infertile women with hormonal imbalance and infertility duration of (5.51 ± 3.21) years. The control group was 21 healthy fertile women with normal hormonal profile. The study was approved by the ethic committee of Tripoli University **(Ref: BEC-BTRC 03-2018)**. The study was fully explained to each participant and informed consent was sought from all participants. A short questionnaire was used to collect detailed information including reproductive hormonal levels and medical diagnosis. Patients enrolled according to the following inclusion criteria: absence of any chronic diseases such as diabetes, high blood pressure, etc., or any sexual diseases, negative serological tests for hepatitis B virus, hepatitis C virus and human immunodeficiency virus. In addition, patients under treatment with any medication were excluded.

### Blood Sample Collection and Processing

Blood samples were obtained from February to April 2019. A sample of 5 ml venous blood were drawn using sterile disposable plastic syringes on day 2 of menstrual cycle of patients and controls. Four ml of blood were collected in plane tube and after centrifugation, sera samples were immediately separated and stored at (−80 °C) for batch analysis. One ml of blood was collected in tube containing 20 µl heparin for blood smear. The heparinized blood was processed within 2 hours after venipuncture (Björksténm, 1974).

### ROS Assessment by the novel NBT Method

The NBT dye 0.1 % was prepared by dissolving 10 mg of NBT powder (Sigma Aldrich) in 100 ml of phosphate buffered saline (PBS, pH = 7.2) and mixed at 37°C for one hour.

### Determination of Superoxide Reactivity in Blood Neutrophils

Equal amounts of 10 µL of each NBT buffer solution and fresh heparinized blood were mixed gently for approximately 30 seconds then incubated at 37°C for 15 min. The samples were left standing at room temperature for 15 min, then 5 µl were placed on a clean glass slide and a thin blood smear was made. The blood smears were dried and fixed using methanol 95% for 5 min (Adewoyin and Nwogoh, 2014).

Smears were stained using Wright’s stain for 30 min, mounted in DePex and examined microscopically according to Battlement’s method (Vilchez, 2020). Oil immersion objective (100x) was used for neutrophil counts with intracytoplasmic formazan deposit (Humbert *et al*., 1973).

### Determination of Superoxide Reactivity Level in Serum

In principle, the assay is based on the production of a colorimetric reaction that gives rise to stable formazan by incubating 20 µl of nitro-blue tetrazolium (NBT) solution with 20 µl of serum for 2 hours at 37°C. Quantification of the changes were performed by visual observation, using a color scale that ranged from a light pink to a dark purple depending on the concentration of superoxide anion and NBT reduction (Gosálvez et al, 2017; **Figure 1)**.

**Figure 1:**
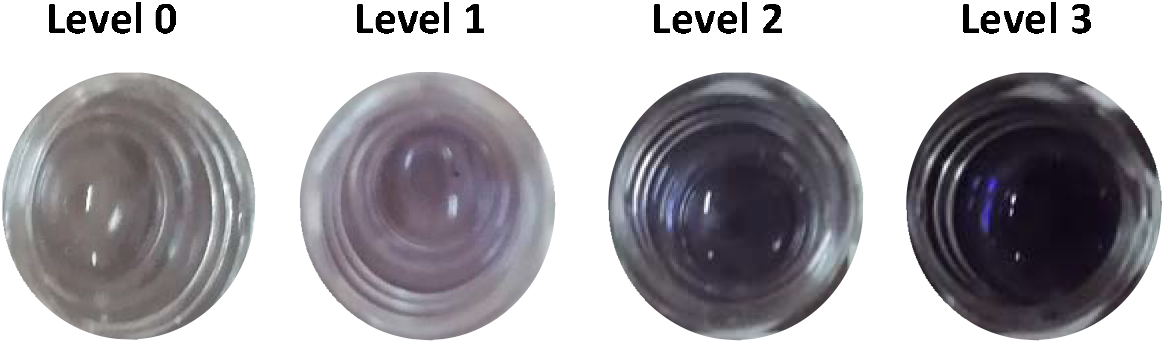
NBT reduction Colorimetric Levels. The color scale was classified to four levels: level 0 = no detectable reactivity, level 1=low reactivity, level 2=medium reactivity, Level 3= high reactivity.

### Determination of Malondialdehyde

Serum MDA levels were measured based on the thiobarbituric acid reactive substances (TBARS) method using an MDA assay kit (ab118970; Sigma Aldrich, UK), in which the MDA-TBA adduct in the serum was generated due to interaction between thiobarbituric acid TBA and MDA. A 20 μl of serum sample was gently mixed with 500 μl of 42 mM sulphoric acid and 125 μl of phosphotungstic acid in a microcentrifuge tube. Samples were then incubated for 5 minutes at room temperature, and centrifuged at 1300 rpm for 3 minutes. The pellet was collected and resuspended in 200 μl ddH_2_O and 2 μl of butylated hydroxytoluin (BHT) (100X) on ice. A TBA reagent (600 μl) were added into the standards and samples vials. The adduct MDA-TBA generated after incubation at 95◦C for 60 minutes and cooling at room temperature in an ice bath for 10 minutes and measured at 532 nm using a microplate reader.

Calculation: The concentration of MDA in the samples was calculated as:

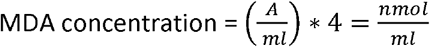

A= amount of MDA in sample calculated from the standard curve (nmol).

ml = original serum volume used.

4 = dilution correction for using 200 μl of the 800 μl reaction mix.

### Statistical Analysis

Statistical analysis were performed using Statistical Package for the Social Sciences software (IBM SPSS statistics 22.0, Chicago, USA). Data were not normally distributed as assessed by Shapiro-Wilk test, so results were expressed as median and interquartile range (IQR). Mann-Whitney test was used to assess differences between the infertile and fertile groups. Correlation analysis was measured using Spearman’s correlation coefficient test. *P-values of* < 0.05 were considered statistically significant.

## RESULTS

### Lipid peroxidation

Oxidative stress as indicated by the lipid peroxidation marker was significant increase in the serum samples of infertile patients compared to controls; MDA concentration in infertile women was (median = 5.6; IQR = 11.00-0.800) compared to controls (median = 1.7; IQR = 2.20-0.00; P < 0.0001; **Figure 2)**.

**Figure 2:**
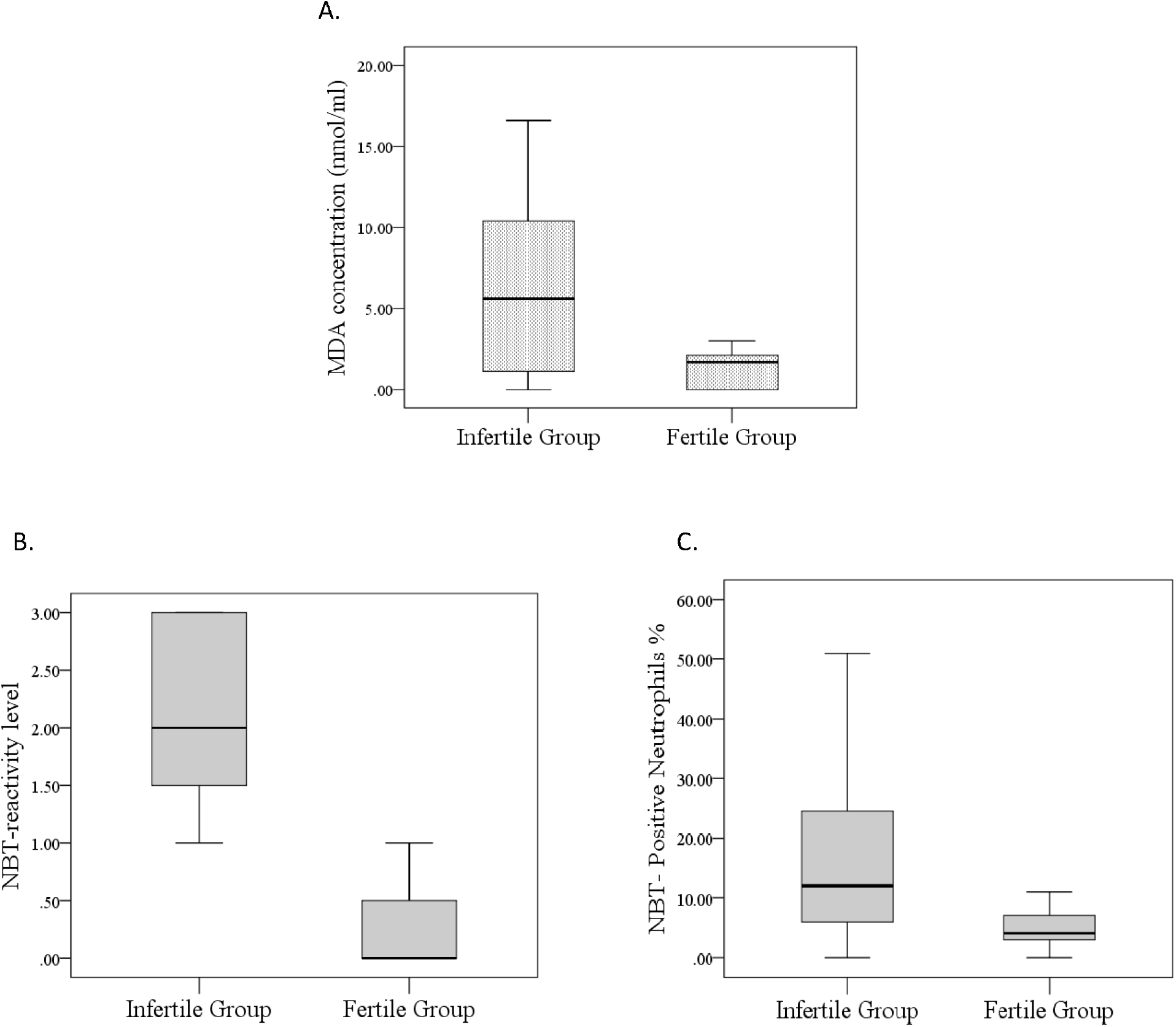
Comparison between infertile and fertile groups. Results are shown as Box and Whisker plot. The bottom and top of the ‘box’ represent the 25th and 75th centile, respectively, while the line within the box represents the median value. The ‘whiskers’ represent the range. A: MDA, B:NBT-Reactivity level and C: NBT-Positive Neutrophils. * p < 0.05, ** p < 0.01, *** p < 0.001, are considered statistically significant.

### NBT-Reactivity level

The presence of oxidative stress was also confirmed by colorimetric scale levels using the original NBT method, where NBT is reduced by superoxide anion, which reflects the concentration of superoxide anion in the serum of infertile and fertile subjects (median = 2; IQR = 3-1) and (median = 0; IQR = 1-0) respectively, P < 0.0001; **Figure 2)**.

### NBT-Positive Neutrophils

The neutrophils also showed positive NBT-reaction for the infertile group (median = 12; IQR = 25-6) compared to their corresponding fertile group (median = 4; IQR = 7.25-2.75; p < 0.0001; **Figure 2)**. The microscopic examination showed that the NBT reaction with superoxide anion was significantly elevated within neutrophils from the patients group compared to controls. The NBT-positive neutrophils had dark purple formazan deposits (toxic granulation) in the cytoplasm comparing with normal granulated neutrophils. Nevertheless, eosinophils, basophils lymphocytes, monocytes and red blood cells did not shown any reaction with NBT dye **(Figure 3)**.

**Figure 3:**
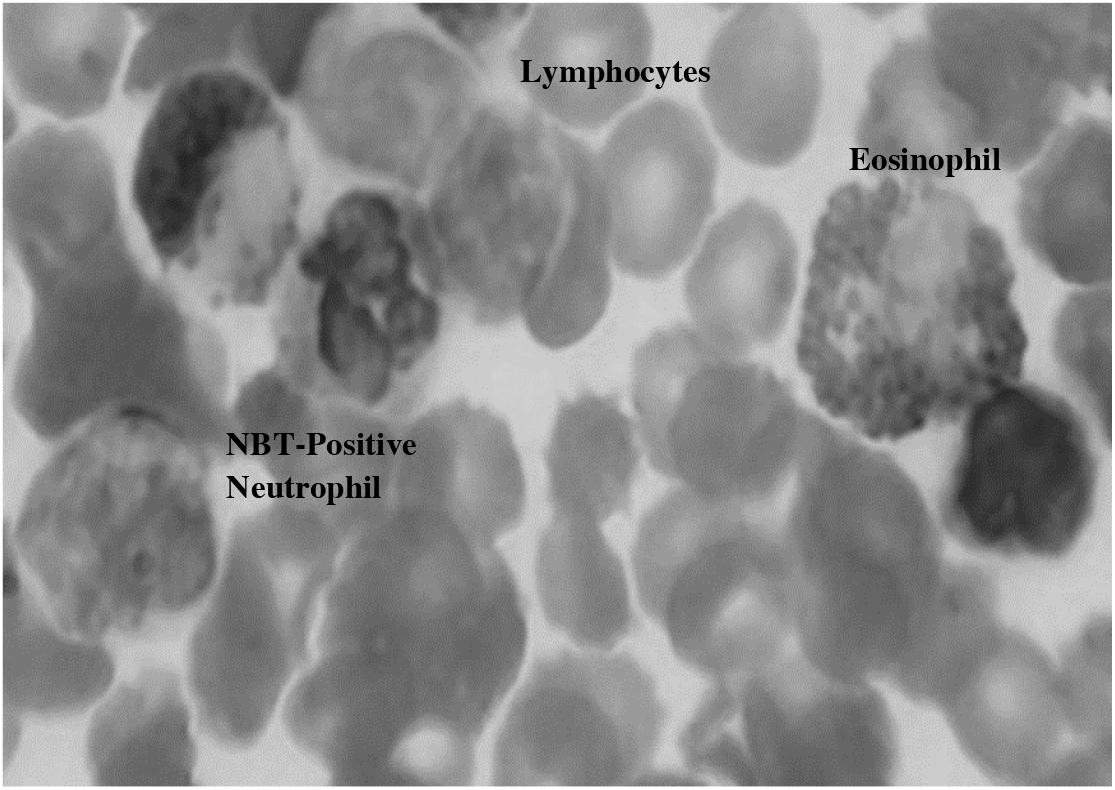
Visualization of Blood Cells. NBT-Positive Neutrophils in comparison with normal Lymphocytes and Eosinophils (NBT and Wright’s-1000X).

### The Correlation Between Oxidative Stress-Related Parameters

The Spearman’s test illustrated a weak negative but significant correlation between increasing serum MDA and decreasing TAC levels (r = −0.243, p = 0.012), while MDA correlated positively with serum NBT-reactivity level (r = 0.410, p = 0.000) and NBT-positive neutrophils (r = 0.207, p = 0.047). Furthermore, there was a significant correlation between NBT-reactivity levels and NBT-positive neutrophils (r = 0.510, p = 0.000; **Figure 4)**. The NBT-positive neutrophils was significantly correlated with age (r = 0.216, p = 0.023) as well as NBT-reactivity level (r = 0.274, p = 0.002; **Figure 5)**.

**Figure 4:**
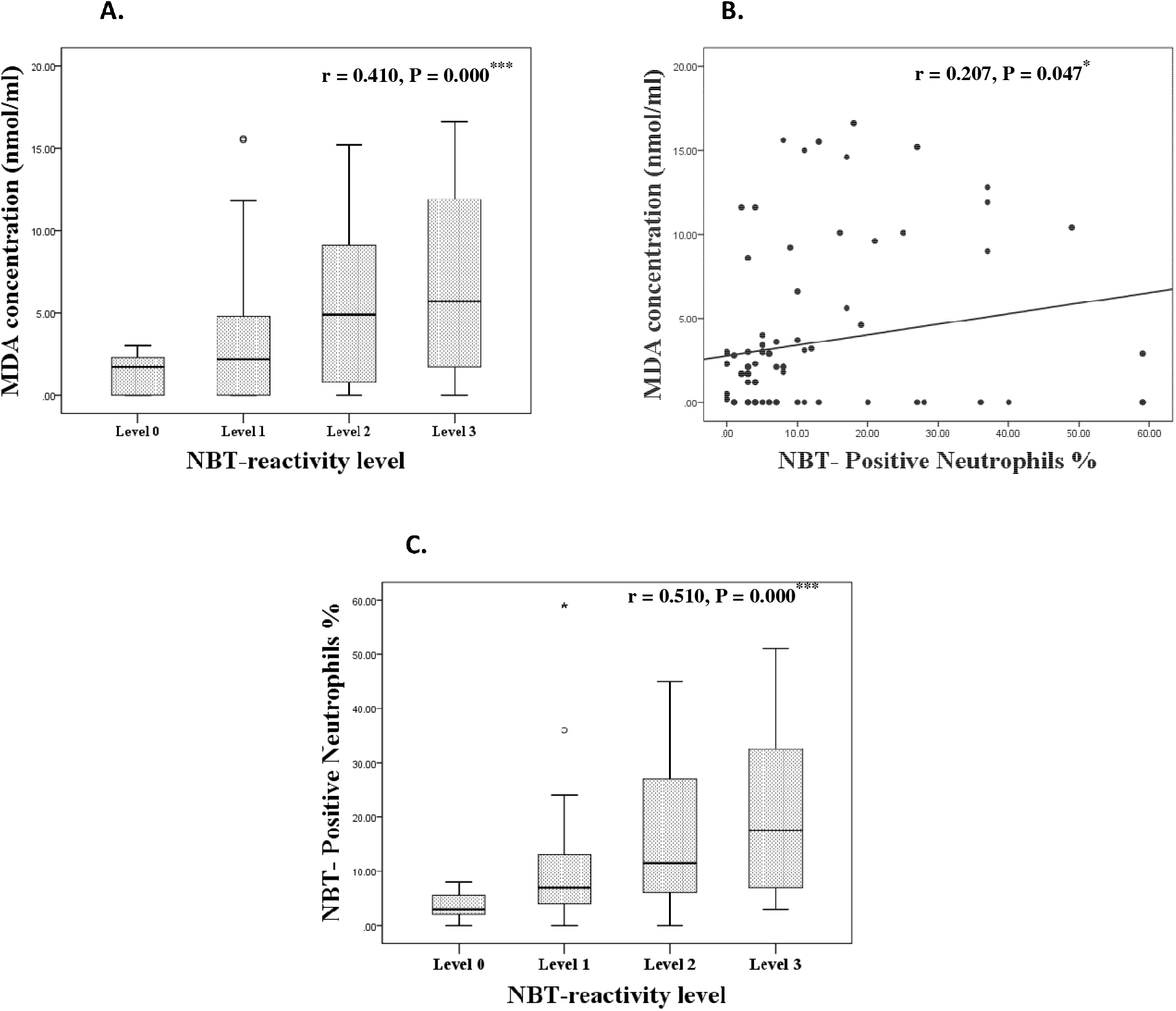
Correlation Between Oxidative Stress Parameters. A: MDA and NBT-reactivity level, B: MDA and NBT-Positive Neutrophils. C: NBT-positive Neutrophils and NBT-reactivity level. *** p < 0.001, ** p < 0.01, * p < 0.05 are considered statistically significant.

**Figure 5:**
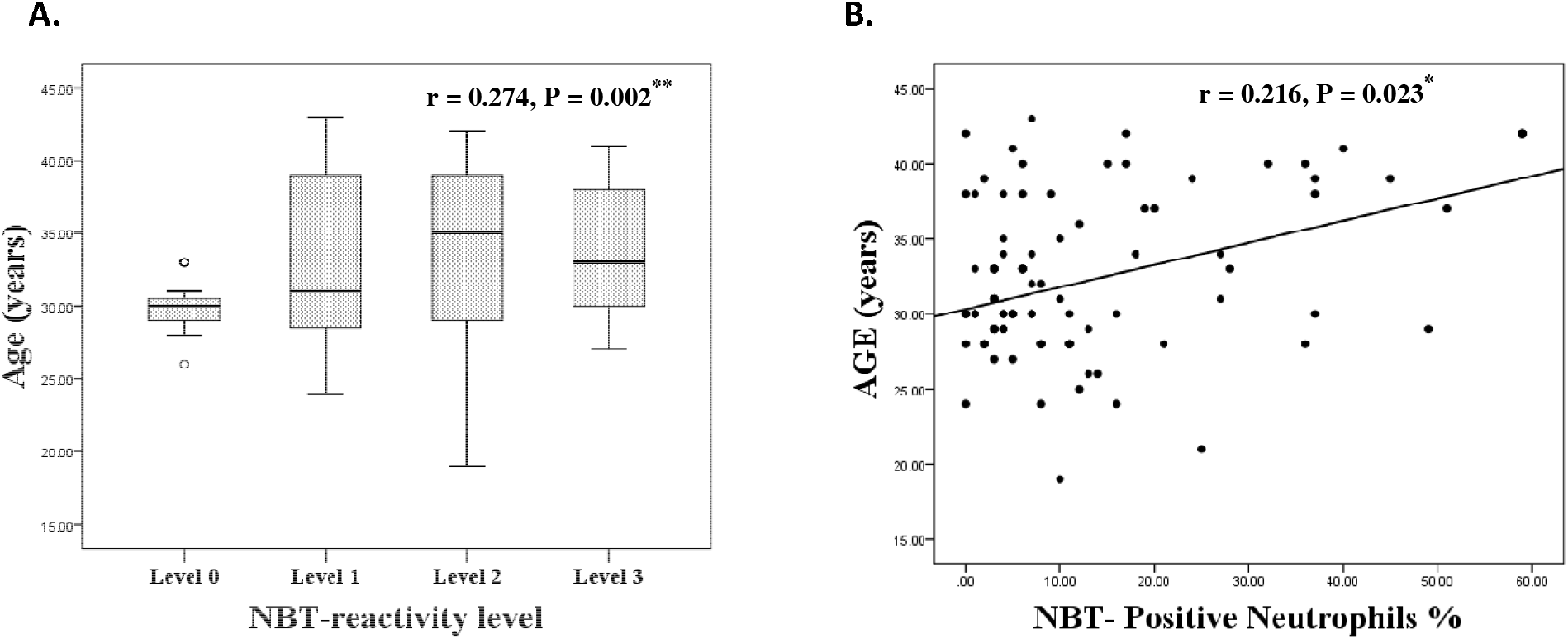
Correlation Between the Investigated Oxidative Stress Parameters and age. A. NBT-Reactivity Level and age. B: NBT-Positive Neutrophils% and age. * p< 0.05 are considered statistically significant.

## DISCUSSION

Infertility is a common problem experienced by many women at reproductive age (Isong *et al*., 2017; Palani, 2016). A large body of evidence suggests that oxidative stress is closely associated with several pathological conditions, where increased oxidants impair the antioxidant defense system and, may over time, contribute to female infertility and poor pregnancy outcomes (Agarwal *et al*., 2012). A major strength of our study is the development of a novel NBT method to assess biomarkers of oxidative stress in infertile women.

Recent work revealed the signs of oxidative stress in infertile women as indicated by increased lipid peroxidation levels and superoxide anion overproduction in the blood samples of infertile women, which indicated an imbalance of redox homeostasis **(Figure 2)**. The most important correlation found was a significant positive correlation between NBT-reactivity level and NBT-Positive neutrophil, MDA and each of NBT-reactivity level and NBT-Positive neutrophil. Furthermore, age was positively correlated with NBT-reactivity level and NBT-Positive neutrophil. Previous studies showed that the peroxidative damage is increased in infertile women which reduces the possibility of conception (Agarwal, *et al*., 2012). Our results demonstrated that MDA levels were higher in infertile women compared to controls, which is in agreement with (Ruder *et al*., 2008; Veena *et al*., 2008; Rad *et al*., 2015; Palani 2016; Becatti *et al*., 2018; Sivaharini *et al*., 2018).

It has been suggested that ovarian dysfunction and polycystic ovary syndrome (Cahill & Wardle, 2002), which results from higher total oxidant status (Cimino *et al*., 2016; Lu *et al*., 2018). Therefore, oxidative stress is not only associated with reproductive disorders, but also has the potential risk of causing infertility in women (Prasad *et al*., 2015). Earlier studies have found that oxidation of lipids in the follicles has deleterious effects on oocyte. This causes peroxidation of oocyte membrane lipid and loss of membrane viability which lead to premature oocyte death and infertility (Dunning & Robker, 2012; Wathes *et al*., 2007). Our results demonstrated that increased MDA levels are correlated significantly with increased NBT-reactivity levels (Ogunro *et al*., 2014; Palani, 2016), suggesting that ROS production can enhance gene expression, which induces lipid peroxidation and MDA formation.

Peripheral leukocytes reflect the condition of the whole organism and thus are a valuable model to study the pathophysiology of oxidative stress-related disorders (Appasamy *et al*., 2008). Leukocytes can show the homeostasis variation due to oxidative stress, which can be responsible for cell damage. Phagocytes are one of the principal sources of ROS, which contain NOX that generates large amounts of superoxide anion and hydrogen peroxide on the outer surface of the plasma membrane (Ahmad *et al*., 2017). Becatti *et al*. (2018) reported that in infertile women, oxidative stress is supported by enhanced ROS generation including superoxide anion in leukocytes compared to controls. The present study demonstrates an increase in ROS, as indicated by high percentage of NBT-positive neutrophils and serum NBT-reactivity levels. Both biomarkers reflect increased superoxide anion generation. Interestingly, we observed a positive and significant correlation between both parameters. This suggests that neutrophils are a contributing factors of increased superoxide anion and MDA levels.

Neutrophils are an important source of ROS in Grraffian follicles. Thus, it has been suggested that ovarian steroidogenesis can induce neutrophils respiratory burst (Rodenas *et al*., 2017), subsequently increase ROS production, especially superoxide anion (Ahmad *et al*., 2017). In agreement with these reports, our results showed a significant correlation between MDA levels and the percentage of NBT-positive neutrophils. These showed that excessive ROS have a negative impact on ovarian functions, which are an important predictor of women infertility. In addition, these results showed indirectly the weakness and reduced capacity of antioxidant system to neutralise oxidants and free radicals. Subsequently lowers the probability of conception.

Ovulation is commonly identified as an inflammatory process. It is induced by biochemical and cellular changes leading to mature ovum release. Subsequently, it upregulates specific genes such as prostaglandin synthase (II), which is associated with ROS production (Shkolnik *et al*., 2011). This suggests that problems associated with ovulation might be linked to ROS overproduction by follicular cells including neutrophils.

Age is a major risk factor for women’s infertility. Several studies showed a strong relationship between age and infertility (Prasad et al., 2015) and that oocyte quality decreases with an increase in maternal age, due to mitochondrial changes resulting from excessive ROS production (Agarwal et al., 2012; Ahmad et al., 2017). Menstrual blood is a way through which women lose iron, which potentiates oxidative damage, and decreases estrogen production along with its antioxidant properties. Our study showed that superoxide production by neutrophils and its level in serum increased with women’s age. Taken together, these results suggest that increased age may predispose women to oxidative stress-related diseases (Agarwal and Allamaneni, 2004).

## CONCLUSION

In this study we described a novel NBT method for detecting the presence of ROS in blood neutrophils and serum. It could be used to assess oxidative stress status in infertile women. Thus, the NBT method could potentially be a very useful diagnosis tool for women’s infertility and may play a role in developing appropriate management strategies to support infertile women.

## ACKNOWLEDGMENTS

The authors would like to acknowledge the support of Dr Muna Ben Salah Dr. Abdelkareem El Fallah, Tripoli Center for Infertility Treatment Head of Haematology Department at Metiqa Hospital, Triopoly, Libya.

